# Characterising the microbial and antimicrobial resistance signatures of hospital-acquired pneumonia using nanopore metagenomic sequencing

**DOI:** 10.1101/2025.08.09.669460

**Authors:** Cedric CS Tan, Alp Aydin, Dewi R Owen, Sylvia Rofael, David Brealey, Mark Peters, Themoula Charalampous, John R Hurst, Timothy D. McHugh, David M. Livermore, Vanya Gant, Justin O’Grady, Francois Balloux, Lucy van Dorp, Virve I Enne

## Abstract

Hospital-acquired pneumonia (HAP) is a significant burden in nosocomial settings, yet its microbial underpinnings remain poorly understood. Here, we leverage shotgun nanopore sequencing to characterise the respiratory microbiomes of 250 HAP patients in a UK multi-site cohort, validating these using paired PCR and culture assays. Sequencing identified the dominant microbes implicated in HAP, including detection of probable pathogens in 49 PCR- and culture-negative cases. We found a high prevalence of fungi in 81/239 (34%) in HAP patients, of whom 26/81 (32%) were PCR/culture-negative, suggesting that fungi may represent an under-investigated component of HAP, whether as colonists or pathogens. Although HAP is clinically sub-categorised based on the use and duration of ventilation before disease onset, we found that the microbial profiles of these sub-groups were indistinguishable. We also found a concerningly high proportion of multi-drug-resistant microbes in HAP patients, with 21% of assembled bacterial genomes harbouring acquired antimicrobial resistance (AMR) genes that confer resistance to at least three classes of antimicrobials. This included high AMR gene carriage associated to *Staphylococcus epidermidis*, which may be an important reservoir of AMR, though typically viewed as a commensal. Our work provides extensive metagenomic characterisation of HAP, underscores the value of metagenomics in describing its complex aetiology, and further prompts its potential role for pathogen detection, resistance profiling and treatment.

## Introduction

Hospital-acquired pneumonia (HAP), defined as pneumonia that occurs 48 hours or more after hospital admission, is one of the most common healthcare-associated infections (HAI), accounting for approximately 15-20% of all HAIs^1^. HAP exacts a sizeable economic burden, estimated to cost the US healthcare system >$3 billion per year^2^. Around 40% of HAP cases present as ventilator-associated pneumonia (VAP), which is defined as a pneumonia occurring 48 hours after initiation of invasive mechanical ventilation^3^. The remaining cases can be further categorised as ventilated HAPs (V-HAP), for patients who are ventilated but developed pneumonia prior to or less than 48 hours after ventilation, and non-ventilator-associated HAPs (NV-HAP). HAP is associated with extended length-of-stay, prolonged mechanical ventilation, and high mortality in vulnerable patients^4–7^, with poorer outcomes for the immunosuppressed or when the causal infectious agent is multi-drug resistant.

The prevailing recommendation for the aetiological diagnosis of HAP is microbial culture of lower respiratory tract specimens^5,8,9^ such as bronchoalveolar lavage (BAL) or endotracheal aspirates (ETAs). Thus, most studies describe the microbial aetiology of HAP using culture-based approaches^10–16^. However, culture is slow and insensitive, yielding no identifiable pathogen in 30-50% of cases^17^, with pathogen detection sometimes compromised owing to antimicrobial administration prior to sampling. This results in a significant underestimation of the microbial diversity responsible for HAP and biased prevalence estimates. Moreover, investigations typically focus on defining a single aerobic bacterial pathogen, without considering the possibility of a more complex aetiology. Identifying putative infectious agents is challenging since the lung, historically thought to be sterile, is now widely accepted to harbour its own microbiome^18^ and the shift that occurs as this healthy lung microbiome transforms to that of HAP and VAP is still poorly understood.

Understanding the microbial component and evolution of HAP is, however, key to developing more effective clinical management and improving patient outcomes. For example, it has been suggested that nearly half of HAP cases involve polymicrobial infections (i.e., caused by more than one putative pathogen)^19^ and that these are associated with a lower 30-day mortality rate than for monomicrobial infections^20,21^. Estimating the prevalence of polymicrobial infections, and the microbes involved, consequently may aid patient risk stratification.

Advances in metagenomic next-generation sequencing (mNGS) technologies have paved the way for culture-independent analysis of patient samples, with significantly faster turnaround times and superior sensitivity for pathogen identification, using both non-invasive and invasive clinical samples^22–24^. They indicate that HAP patients harbour a distinct microbial community signature characterised by low within-sample species diversity (alpha diversity), and a lower abundance of non-pathogenic *Streptococcus* and other commensal species, when compared with non-HAP controls^25^. Leveraging these signatures may enable development of novel diagnostic and prognostic approaches to reduce the health burden of HAP. In addition, metagenomics can enable the characterisation of antimicrobial resistance (AMR) profiles^22^ or virulence factors associated with disease pathogenesis and patient outcomes. Finally, high-coverage complete bacterial genomes can sometimes be generated within two hours of sequencing^22^, potentially allowing for rapid establishment of the routes of transmission during a disease outbreak.

In this cross-sectional, multi-site UK study, we employed nanopore metagenomic sequencing to characterise the microbial component of the different HAP subgroups. The patient samples studied represent a subset of the INHALE WP1 study,^26^ which primarily compared the results of two commercial respiratory multiplex-PCR panels with those of microbiological culture for 652 HAP/VAP patients at 15 ICUs in England. Nanopore metagenomic sequencing was undertaken on the samples from 250 of these patients. We examined the microbial diversity present in these samples and the prevalence of pathogens putatively associated with infections. We additionally investigated the differences in microbial aetiologies and AMR resistance profiles for the different HAP subgroups. We further assembled genomes of the dominant pathogens involved to explore the molecular epidemiology of these infections for the samples with a sufficient yield of microbial sequencing reads.

Our results indicate no consistent differences between the microbial communities of HAP versus VAP patients. HAP and VAP patients did, however, differ in their carriage of antimicrobial resistance genes, which may reflect differences in the antimicrobial prescribing practices typical for these clinically defined categories of patients. We additionally identified a wide diversity of potentially causative bacterial pathogens, and noted the presence of fungi, whether as colonists or contributory agents, in a third of HAP patients. Overall, our results shed light on the diversity of microbial taxa and their AMR profiles in HAP, demonstrating the power of metagenomic sequencing in understanding the aetiology and epidemiology of hospital-acquired infections.

## Results

### Robust identification of pneumonia-associated microbes

We investigated non-invasive and invasively sampled specimens from our primary cohort of 250 critically ill HAP patients recruited from seven ICUs in the UK (54 NV-HAP, 49 V-HAP, 147 VAP; **Extended Data Fig. 1**). To characterise the microbial diversity associated with the different HAP subgroups, we processed these samples using an established host-depletion protocol designed to enrich microbial DNA for shotgun nanopore metagenomic sequencing^22^, with the inclusion of negative sequencing controls to account for possible laboratory contamination. We subsequently performed taxonomic assignment of sequencing reads using Kraken2^27^ and applied a set of bioinformatic filters – minimising the effects of artefactual signals arising due to sequencing analysis errors, index hopping and laboratory contamination (see **Methods**) – to identify the presence of microbial taxa. A detailed assessment of the effectiveness of these filters are provided in **Supplementary Note 1**. We compared the species detected through this metagenomic sequencing with those species found by two commercial multiplex-PCR panels (BioFire FilmArray Pneumonia Panel and Curetis Unyvero) and microbial culture, as published previously^26^. All filtered taxonomic assignments are provided in **Supplementary Table 1**.

Of the 250 patient samples we sequenced, 239 (96%) yielded information on microbial content, defined by having at least 100 reads which could be assigned to microbial taxa. Following bioinformatic filtering, we detected a median of five microbial species per sample, though with up to 38 microbial species identified in one case. Since the distinction between pathogens and commensals is sometimes unclear, particularly in non-invasive samples liable to contamination by the flora of the upper airways (or at sampling), it is likely that only a subset of these microbes was associated with the clinical pneumonia. Since the PCR and (in general) culture assays employed here as comparators report only microbes with established clinical associations to pneumonia, we loosely categorised microbial species as ‘pneumonia associated,’ or not, based on whether they are sought and conventionally reported by either of these methods. The list of pneumonia-associated species defined on this basis is provided as **Supplementary Table 2**.

The most common pneumonia-associated species identified via sequencing were *Staphylococcus aureus* (n=55, 22%), *Klebsiella pneumoniae* (n=37, 23%), *Pseudomonas aeruginosa* (n=29, 12%), and *Escherichia coli* (n=25, 10%) (**Fig. 1a**). Additionally, we found a considerable number of polymicrobial infections (n=66, 28%), defined by the presence of more than one pneumonia-associated genus. For instance, we identified one case of up to five pneumonia-associated genera detected in a single patient (**Fig. 1b**). The prevalence of pneumonia-associated taxa and of putatively polymicrobial infections detected via sequencing were largely concordant with that estimated via PCR and culture (**Extended Data Fig. 2a and b**, respectively). These findings indicate that metagenomic sequencing can robustly identify pneumonia-associated pathogens.

**Figure 1.**
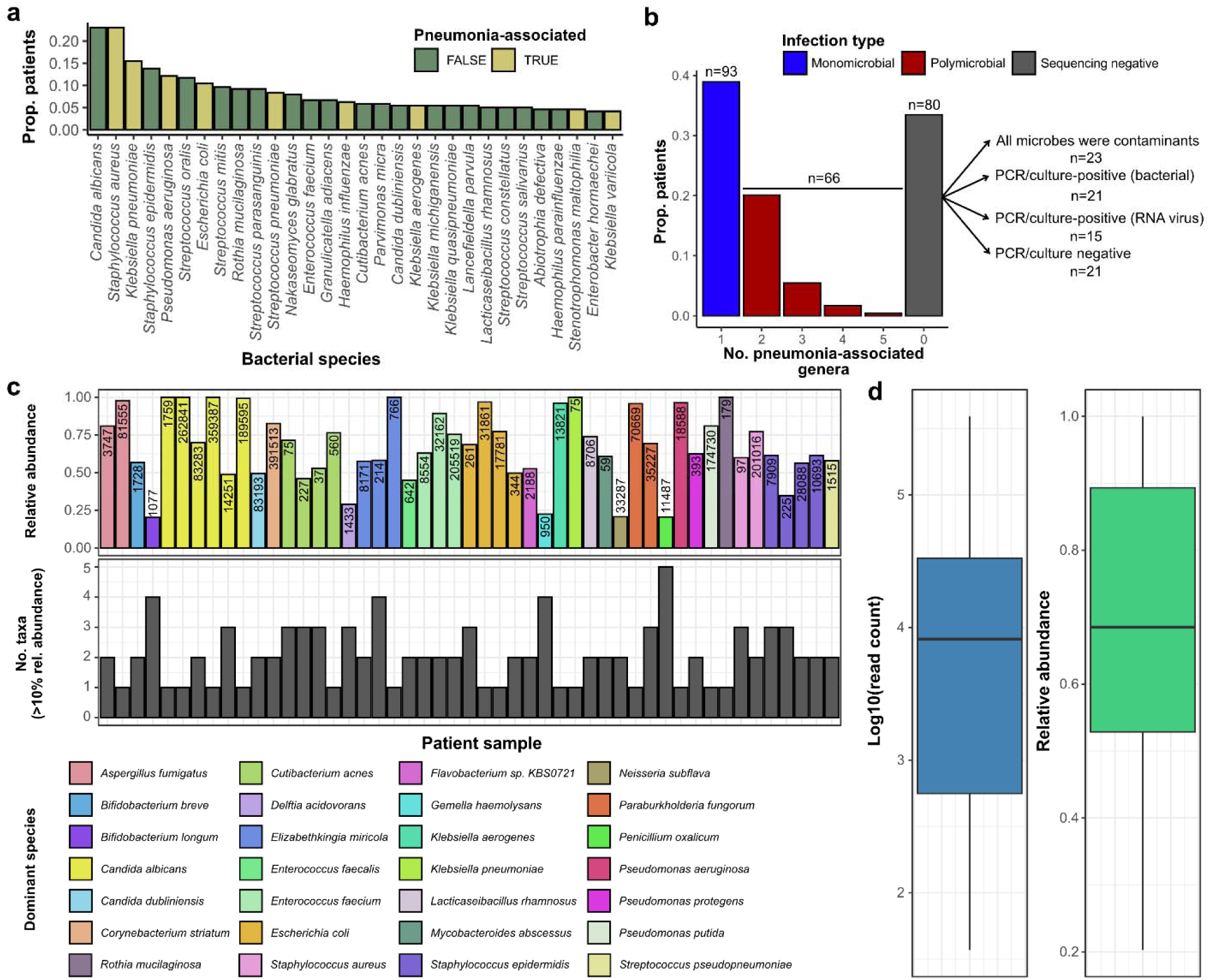
Robust identification of pneumonia associated microbes using Nanopore sequencing. (a) Proportion of patients testing positive for each of the 30 most common microbial species detected through shotgun nanopore sequencing. (b) Proportion of samples classified as mono- and polymicrobial as identified via sequencing (n=239). Polymicrobial infections are defined by the detection of more than one pneumonia-associated genus by sequencing. The breakdown of patient samples with no pneumonia-associated species is also given. (c) Most abundant microbial species detected in the 49 patient samples with no pneumonia-associated pathogens identified via PCR or culture. The number of species present at >10% relative abundance in each sample is also shown. (d) Distribution of read counts and relative abundance of the dominant species identified in (c). Pneumonia-associated species/genera are defined as those ordinarily reported by PCR and/or culture (see main text and **Supplementary Table 2**).

No pneumonia-associated species were identified via metagenomic sequencing in 80/239 (33%) of patient samples (**Fig. 1b; Supplementary Table 3**). For 23 of these, all microbes detected were discounted as they were deemed likely to be associated with laboratory contamination (see **Methods**). Fifteen patients tested positive for RNA viruses via PCR, indicating that these patients may have had pneumonias caused by RNA viruses. These would not have been identified via sequencing, as the protocol we used does not target RNA. It should also be noted that our sampling predated the SARS-CoV-2 pandemic. Twenty-one samples were both PCR and culture negative. The remaining twenty-one were PCR or culture positive for bacterial pathogens, indicating some discordance between sequencing and non-sequencing-based methods, which could have been caused by false positives in the PCR/culture assays, or false negatives in sequencing, though it is difficult to differentiate the relative impacts of either. Notably, the PCR and culture assays were also discordant in 13 of these 21 sequencing negative, but PCR/culture-positive samples, highlighting variation in sensitivity and specificity across all methods.

Of the 239 samples, 50 were PCR/culture negative (**Supplementary Table 3**). Sequencing reliably identified at least one microbial species in 49 of these samples. The most abundant microbial species (henceforth ‘dominant species’) found in these 50 cases included both pneumonia-associated and non-pneumonia-associated microbes, with a tendency towards communities being more polymicrobial in nature (34/49 cases with detection of more than one pneumonia-associated genus) (**Fig. 1c**). Across these dominant species, the median read count was 8171 with a mean relative abundance of 69% (**Fig. 1d**), which is consistent with them exhibiting rapid growth during infection and subsequently dominating the recovered sequencing reads. Additionally, these species had at least a 2.8-fold greater relative abundance in patient samples than in the negative sequencing controls, and passed the set of filters we implemented to account for possible contamination (see **Methods, Supplementary Note 1**). Further, five of the 14 dominant species identified in these samples, assessed as having at least 10,000 Kraken2-assigned reads, showed sequencing coverage biases characteristic of growing bacterial populations^28,29^ (see **Methods**; **Extended Data** Figure 3). These findings demonstrate that metagenomics holds promise to identify pneumonia-associated microbes in cases where other assays were unsuccessful.

### Fungi are frequently observed in hospital-acquired pneumonia

Pneumonia can be caused by diverse microorganisms, mostly thought to comprise bacteria and viruses. With the exception of *Aspergillus*, fungi detected in the respiratory tract are often considered to be colonists that overgrow in the upper airways as a result of antibiotic-induced disruption of the respiratory microbiome^30^. Strikingly, we recovered evidence of the presence of fungal genera in 81 (34%) of the 239 HAP patients whose samples were successfully sequenced. Of these, 57 (24%) harboured fungal taxa at a more stringent detection threshold of >5% relative abundance. Moreover, *Candida*, *Aspergillus* and *Nakaseomyces* were found at more than 50% relative abundance in 29/81 (36%) of these fungi-positive patients (Fig. 2a), and 26/81 (32%) of these had no other microbes identified via PCR or culture. None of these fungal taxa were found in negative sequencing controls, indicating that laboratory contamination is unlikely. To confirm that the identification of fungal genera were not a result of taxonomic misclassification, we performed a more-stringent read mapping analysis using Minimap2^31^ for *Candida*, *Aspergillus* and *Nakaseomyces* species. We found a near perfect correlation (*r*=0.999) between the number of reads mapped to these species by Minimap2 and Kraken2, indicating that these hits were not taxonomic misassignments. Furthermore, 21/81 (26%) of these samples showed presence of fungal growth in culture, but these observations were not reported initially, in line with frequent culture-reporting practices.

**Figure 2.**
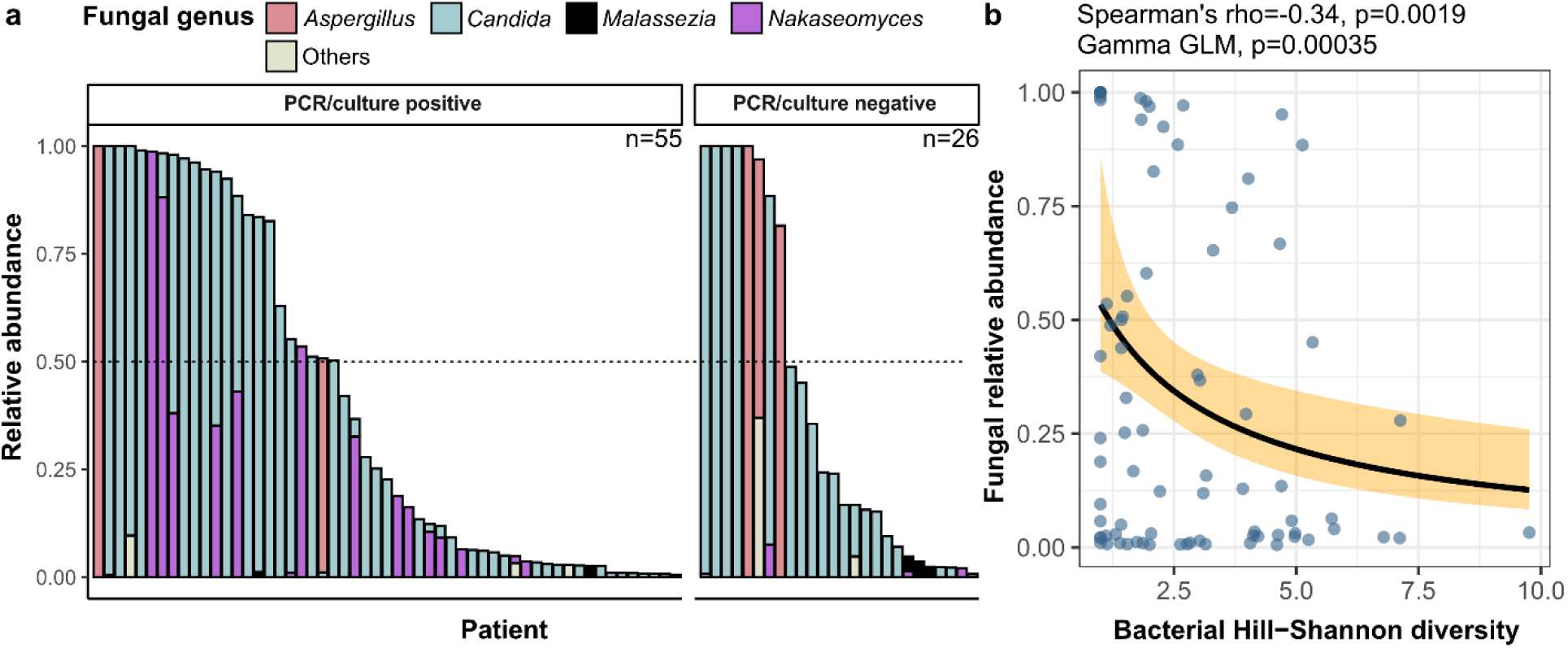
Detection of fungal DNA in hospital-acquired pneumonia. (a) Relative abundance of fungal genera in pneumonia patients (n=81) after accounting for laboratory contaminants and sequence analysis artefacts (see Methods and Supplementary Note 1). Samples are stratified by PCR/culture positivity status. (b) Inverse correlation between the summed relative abundance of fungal genera and Hill-Shannon diversity of bacterial genera. P-values were calculated using a two-sided t-test of the Spearman’s *ρ*, and a two-sided z-test on the parameter estimate in a Gamma generalised linear model correcting for antibiotic exposure and sample type.

In contrast, we were not able to detect evidence of fungal taxa in metagenomic data generated from a separate cohort of eight healthy individuals, who were recruited from primary health care practices as a comparator group in a separate study^32^ and whose samples were processed with the same laboratory protocol. This indicates that it may not be typical to observe these fungal species in the respiratory tract of individuals without pneumonia, in line with previous findings^33,34^.

Notably, we found that the relative abundance of fungi was negatively correlated with bacterial alpha diversity (Spearman’s *ρ*=-0.34, p=0.0019), even after correcting for antibiotic exposure and sample type (Gamma general linear model, two-sided z-test, z=2.92, d.f.=75, p=0.0035) (Fig. 2b). This indicates that the proliferation of these fungal taxa is linked to the potential disruption of the respiratory microbiota, though whether this is a causal association and the direction of causality will require further study. While the conventional wisdom is that fungi, other than *Aspergillus* spp., are likely non-pathogenic colonisers in these patients, our results indicate that their role in the pathogenesis of HAP (including VAP) may warrant re-examination.

### HAP subgroups display similar microbial signatures

HAP, which includes NV-HAP, V-HAP and VAP, is likely caused by the aspiration of pathogenic microbes that have colonised the upper respiratory tract (URT), or contaminated aerosols from environmental sources^19,35^. Pneumonia associated with mechanical ventilation (i.e., V-HAP and VAP) is believed to be accelerated by the insertion of an endotracheal tube, which both disrupt normal mucociliary clearance^36,37^ and provides opportunity for biofilm formation on an avascular structure. Given differences in the likely routes of infection, we explored whether the microbial profiles of the three HAP subgroups differed.

Initially we applied a principal coordinates analysis to the microbial profiles, finding no obvious stratification based on HAP subgroup (Fig. 3a). We then applied multivariate analysis of variance (MANOVA) to correct for the effects of sequencing depth and sampling site to more rigorously test for differences in the principal coordinates (PCo) by HAP subgroup (see **Methods**). We found no significant differences in the diversity of microbial genera by HAP subgroup (PCo1: one-tailed F-test, F=1.45, p=0.238; PCo2: F=1.50, p=0.225). This was consistent across various decontamination thresholds (**Supplementary** Figure 2; **Supplementary Note 1**).

**Figure 3.**
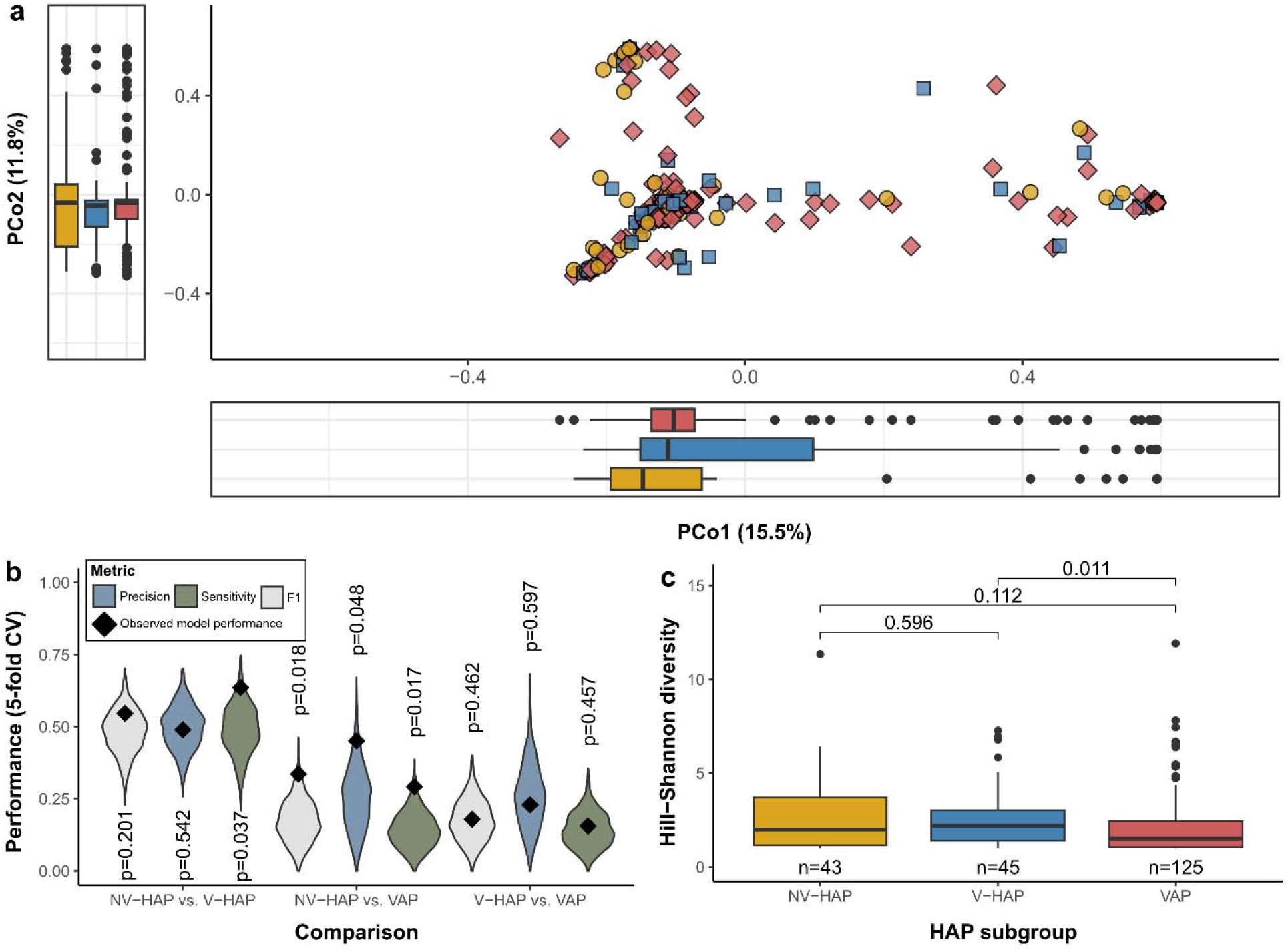
HAP subgroups have similar microbial signatures. (a) Principal coordinates analysis of Bray-Curtis distances, and boxplots showing the lack of clustering in the microbial profiles of NV-HAP (yellow), V-HAP (blue) and VAP (red) samples. (b) Violin plot showing the distributions of average model performance scores for models trained on randomised data. Black diamond points indicate the observed performance scores for models trained on the actual data. P-values were calculated as the proportion of randomisation iterations yielding performance scores at least as high as the observed score. (c) Boxplot of Hill-Shannon diversity stratified by HAP subgroup. Pairwise differences in diversity were tested using two-sided Mann-Whitney U tests and the corresponding p-values are annotated. Boxplot elements are defined as follows: centre line, median; box limits, upper and lower quartiles; whiskers, 1.5-fold interquartile range.

To look for more subtle differences in the microbial profiles of HAP subgroups, we trained gradient-boosted tree classifiers to identify the HAP subgroup of a patient based only on their sequencing-defined microbial profiles. This is a powerful machine learning algorithm that has been successfully used to predict the status of individuals with diseases including sepsis^38^, chronic lung disease^39^ and type II diabetes^40^ based solely on microbial DNA content. To formally assess whether our models were capturing genuine signals in the data, we also compared their performance to that of models trained on randomised data (see **Methods**). We were unable to reliably distinguish between the microbial profiles reconstructed in NV-HAP and V-HAP (precision=0.49, sensitivity=0.62, F1=0.55), NV-HAP and VAP (precision=0.45, sensitivity=0.29, F1=0.34), or V-HAP and VAP (precision=0.23, sensitivity=0.16, F1=0.18) (Fig. 3b). Additionally, the randomisation tests indicated that the models trained to discriminate between NV-HAP and V-HAP, and V-HAP and VAP did not perform better than models trained on randomised data (Fig. 3b). Separately, we tested whether the microbial alpha diversity within samples differed between HAP subgroups, and found subtle differences in the microbial genus diversity (linear regression, F=2.70, p=0.070; Fig. 3c), but not when assessed at the level of the microbial species (F=1.75, p=0.18). Overall, these results suggest that the microbial profiles of different clinically defined HAP subgroups are indistinguishable.

### Antimicrobial resistance profiles of HAP-associated microbes

We next explored the predicted AMR phenotypes of bacteria in our HAP samples. To do so, we predicted the resistance phenotypes of microbes in all 239 samples *in silico* using ResFinder^41^. To minimise the effects of laboratory contamination, we excluded phenotypes of samples that were also identified in their corresponding negative sequencing controls. Across all samples, 64% harboured bacterial genes predicted to confer resistance to at least one antibiotic. The most prevalent included those that confer resistance to commonly used antibiotics, with eight of the 20 conferring resistance to antibiotics recommended by UK and/or US guidelines for the treatment of HAP^42,43^ (Fig. 4a).

**Figure 4.**
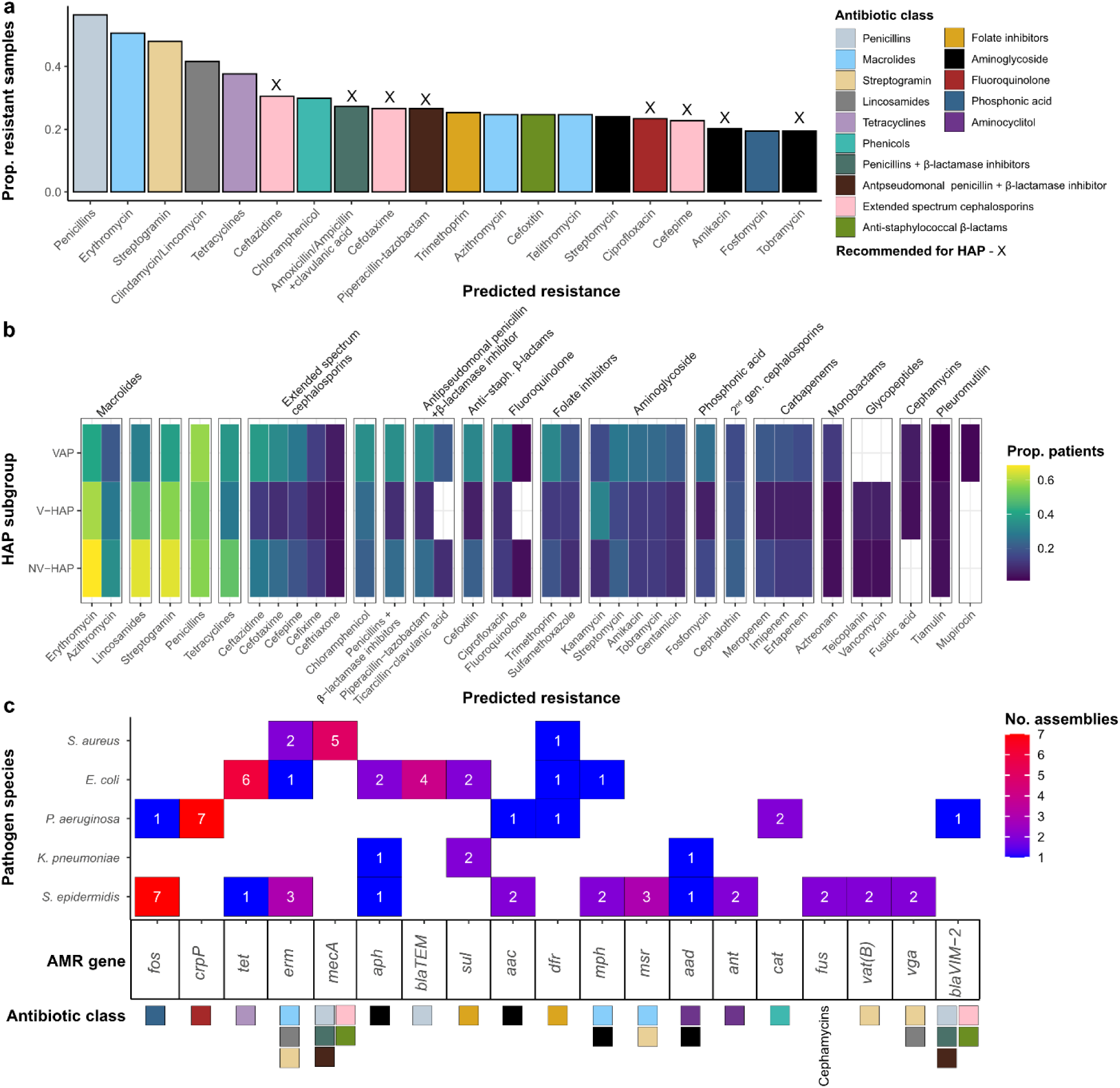
AMR profiles of HAP-associated microbes. (a) Barplot showing the proportion of samples harbouring the 20 most prevalent resistance genes, as detected by ResFinder^41^. The antibiotics recommended for the treatment of HAP by UK/US guidelines^42,43^ are annotated. (b) Heatmap indicating the prevalence of resistance gene carriage to the different antimicrobials stratified by HAP subgroup. Only antimicrobials with at least 10 positive predictions are shown. (c) Heatmap showing the diversity and frequency of acquired AMR genes detected in the top five species with the greatest number of genome assemblies recovered. Resistances intrinsic to a species or genus and mediated by universal chromosomal genes were excluded. The antimicrobial classes that these genes compromise are annotated as in (a).

To test whether the HAP subgroups differed in their AMR gene content, we performed linear regression on a principal coordinates analysis of AMR gene presence (see **Methods**). We were able to detect subtle differences in gene carriage (PCo1: F=2.55, p=0.0817; PCo2: F=3.52, p=0.0323), likely driven by the higher proportion of predicted macrolide, lincosamide and streptogramin b resistance in NV-HAP and V-HAP patients compared to VAP patients (Fig. 4b). Further inspection of the genes associated with these resistance types indicated that 81% were *erm* or *msr* genes (**Extended Data Fig. 4**). These encode ribosomal methylases and efflux pumps, respectively, the former associated with concurrent resistance to macrolides, lincosamides and streptogramin b (MLS_b_)^44^ and the latter to macrolides and streptogramin b.

We then set out to identify the bacterial taxa carrying these AMR genes. To do so we performed *de novo* assembly of the metagenomic sequencing reads to reconstruct draft genome assemblies, which were then subjected to taxonomic classification and screened for AMR gene content, focusing only on genes that were likely to have been acquired, and not those intrinsic to particular species. In total, we recovered 338 assemblies that could be confidently assigned to a bacterial species (see **Methods**), resulting in assemblies ranging from 200kb to 6.4mb in length. The five species for which we recovered the greatest number of assemblies included some typical pneumonia-associated pathogens: *E. coli* (n=42), *S. aureus* (n=42), *P. aeruginosa* (n=24), *K. pneumoniae* (n=22), and *S. epidermidis* (n=16), mirroring our Kraken2 analysis above (Fig. 1a). The most common AMR gene associated with each of these species were *tet, mecA, crpP, sul* and *fos*, respectively (Fig. 4c). Notably, *S. epidermidis*, a common oral and skin commensal, was associated with a large diversity and high frequency of resistance genes, including the *fos* gene, which confers fosfomycin resistance (Fig. 4c). This indicates that *S. epidermidis* may represent a reservoir of AMR genes with potential for horizontal transfer of resistance genes to other organisms, which may be important in the respiratory tract.

Of the 338 contigs, 67 were assessed to be at least 90% complete (median N50=399,066) and of high quality by CheckM2^45^, and spanned 22 bacterial genera and 30 species (**Extended Data Fig. 5**). The bacteria represented by these complete assemblies were predicted to be resistant to between zero to seven antimicrobial classes (**Extended Data Fig. 6**), and 21% were predicted to be multi-drug resistant (defined as resistant to at least three antimicrobial classes that ordinarily are useful against the species), further highlighting the high prevalence of AMR genes in respiratory microbes.

### Molecular epidemiology of pneumonia-associated pathogens

HAP is thought to develop following the aspiration of pathogenic organisms, which may be acquired in the community or within hospital settings. However, aside from documented outbreaks of single pathogens^46,47^, the relative contributions of either mode of acquisition remain unclear. To demonstrate the utility of metagenomic sequencing for characterising the epidemiology of pneumonia-associated pathogens, we assessed the relatedness and sub-lineage affiliations of the complete bacterial assemblies compared with publicly available genomes. We focused on the three species for which we had the highest number of complete assemblies (>=90% CheckM2 completeness): *S. aureus* (n=14), *Haemophilus influenzae* (n=8) and *E. coli* (n=6), and placed these into reconstructed phylogenies of the wider genetic diversity of each species (**Fig. 5a-c**).

**Figure 5.**
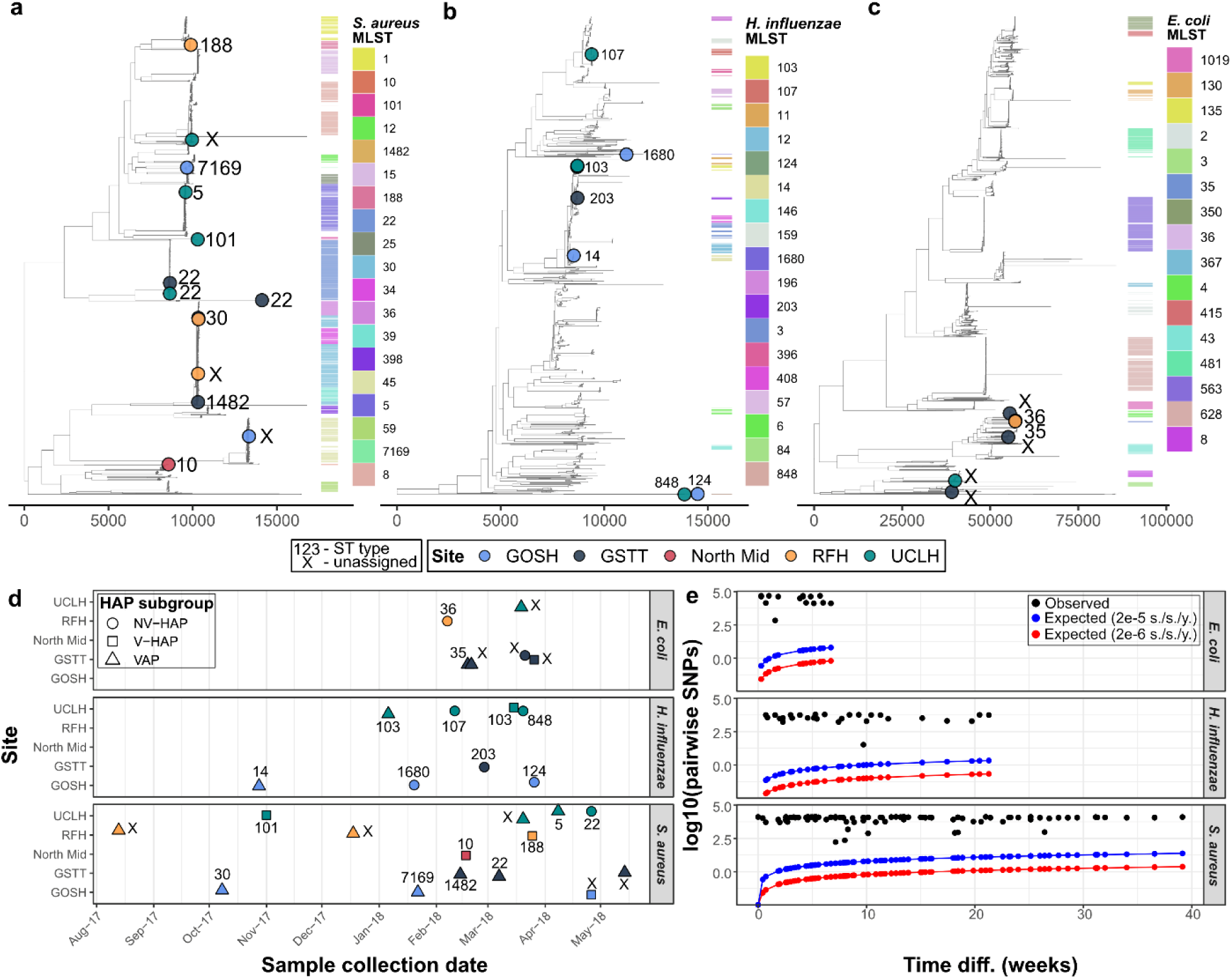
Molecular epidemiology of pneumonia-associated pathogens. Recombination-pruned, core-genome phylogenies of (a) *S. aureus*, (b) *H. influenzae* and (c) *E. coli* assemblies recovered from our samples in the context of publicly available genomic datasets. (d) Distribution of sample collection dates, MLST types inferred based on sequence content, HAP subgroups and hospital sites for the bacterial assemblies recovered from our samples. (e) The observed number of SNPs between all pairwise combinations of recovered assemblies and the differences in sampling dates in weeks. The expected number of SNPs given the previously reported evolutionary rate of 2×10^−6^ substitutions/site/year^48,49^, and given a much higher rate of 2×10^−5^ substitutions/site/year are shown in red and blue, respectively.

For *S. aureus* (**Fig. 5a**) and *H. influenzae* (**Fig. 5b**), we found that our recovered genome assemblies were diverse, phylogenetically interspersed throughout the wider diversity of isolates, and belonging to different Multi-Locus Sequence Typing (MLST) sequence types (**Supplementary Table 4**). This remained the case for assemblies recovered from samples taken from patients at the same hospital, suggesting a wide genetic diversity. For *E. coli*, the assemblies appear clustered but are placed within distinct MLST groups (**Fig. 5c**). There was no obvious pattern in the MLST types of recovered assemblies, regardless of HAP subgroup, within sampling sites, and across time (**Fig. 5d**). To further test whether any of the recovered assemblies could be part of the same transmission chain, we calculated the number of SNPs between all pairs of recovered isolates. Assuming a constant evolutionary rate for genomes within the same transmission chain, the expected number of SNPs between any two assemblies is directly proportional to the difference in their sampling dates. However, the observed number of pairwise SNPs between our assemblies far exceeded the expected number of SNPs given previous rate estimates of ∼2×10^−6^ substitutions/site/year for *E. coli*^48^ and *S. aureus*^49^, and continued to do so even when assuming a 10-fold higher rate (i.e., 2×10^−5^ substitutions/site/year) (**Fig. 5e**). This likely indicates that none of the recovered pathogens were part of the same transmission chain.

## Discussion

In this work, we employed long-read nanopore sequencing to characterise the respiratory microbiomes of 250 HAP patients (54 NV-HAP, 49 V-HAP, 147 VAP) across a UK multi-site cohort. Our metagenomics sequencing workflow, along with assignment and analysis of bacterial and fungal taxa content, provided a detailed characterisation of the microbial composition and AMR profiles associated with these HAP subgroups. We identified a wide range of microbial species, some of which could be characterised at the strain level down to their precise phylogenetic placement. While conventional clinical detection panels tend to focus on a narrow set of causative pathogens, our work suggests that many species may be putatively implicated in HAP, several of which are not routinely sought or reported. While we observe a spectrum in the microbial profiles, our results suggest that the microbial compositions associated with different HAP subgroups are indistinguishable. We further predicted a high prevalence of antimicrobial resistance in HAP-associated microbes. Our work highlights the plethora of clinical, microbiological and epidemiological insights that could be gleaned from shotgun nanopore sequencing from clinical specimens, including those from HAP patients.

In most (49/50) PCR- and culture-negative samples, sequencing was able to detect a diversity of microbes that were not necessarily limited to traditional nosocomial pathogens, some of which exhibited patterns consistent with rapid growth during infection (**Fig. 1c, d**; **Extended Data Fig. 3**). Many of the dominant microbes in these infections have been reported as rare causal agents of pneumonia, including the commensals *Rothia mucilaginosa*^50^*, S. epidermidis*^51^, and *Corynebacterium striatum*^52^ and known human pathogens such as *Mycobacteroides abscessus*^49^, *Aspergillus fumigatus*^5^. However, most of these microbes, when detected by culture, are construed to be colonists or commensals on the basis that they are not commonly considered pathogenic and/or associated with HAP. Our finding aligns with evidence supporting the presence of a microbiome in the healthy lung^18^, which we hypothesise may be disrupted during episodes of pneumonia and mechanical ventilation. We further demonstrate the ability of metagenomic sequencing to provide a relatively unbiased view of the microbiological composition in patient samples, including both aerobes and anaerobes, enabling better characterisation of the microbial aetiology of HAP cases, potentially within clinically-actionable timeframes^22^.

We observed that up to a third of the patients had microbiomes containing fungal taxa, including both yeasts (C*andida, Nakaseomyces)* and mould (*Aspergillus* spp.), with these species sometimes dominating the sequencing reads. Outside of the context of deep immunosuppression, yeasts associated with humans are not recognised to cause tissue-invasive pulmonary disease and most consider them as opportunists, taking over the bacterial niches disrupted by antibiotic therapy. Their presence, and frequent dominance, in the airways of critically ill patients however prompt further consideration for a pathogenic role. Such yeasts trigger highly pro-inflammatory immune responses involving release of cytokines, including but not limited to TNF, IL-1 and IL-6; the local generation of reactive, cytolytic oxygen intermediates; as well as the synthesis of fungal polysaccharides that promotes inflammation through TLR2, TLR-6, NLRP3, and dectin-1^53^. Our frequent detection of *Aspergillus* must be interpreted differently: *Aspergillus* is a constantly-inhaled mould^54^, and its presence may simply reflect inhaled spores; conversely, members of the species have more recently been recognised as a true and often underdiagnosed opportunistic pathogen in ICU patients, particularly among those with suspected VAP^55^. We note that in our dataset we recovered at least eight cases where a fungal species was found to dominate, in the absence of any other pathogen detected via PCR or culture. Our results demonstrating such a high prevalence of fungal sequences prompt further consideration of what consequences their presence has for the patient’s lung pathology.

Despite recovering a broad range of microbial signatures, we were unable to distinguish clinically defined NV-HAP, V-HAP and VAP based solely on analysis of microbial content. This suggests that HAP and VAP may not measurably differ in terms of microbiota or ‘causative agent(s)’. This implies that intubation *per se* may not significantly impact the respiratory microflora; or, at least, it indicates that intubation does not lead to an easily detectable signature once an infection had developed. We did however detect differences in the antimicrobial gene content in HAP versus VAP. In particular, we found a higher proportion of genes conferring MLS_b_ resistance in NV-HAP and V-HAP patients compared with VAP patients. This may reflect our previous observation^26^ of a greater overall broad-spectrum antibiotic burden administered in VAP as opposed to HAP and V-HAP.

Phylogenetic analyses of our recovered bacterial assemblies indicate that, within our dataset, the microbes associated with HAP span a wide diversity of pathogen lineages, and no pair of the sampled patients were likely part of the same transmission chain (**Fig. 5**). We acknowledge that this represents a small sample size. Nevertheless these findings suggest that at least some of the causative microorganism(s) may have been acquired outside the hospital setting, potentially prior to hospital admission. One possibility is that some commonly implicated microbial species reside asymptomatically within patients, who then became susceptible to infection due to critical illness, invasive procedures, or disruption of the respiratory microbiome as a result of antibiotic therapy during their hospital stay. While we lacked the intensity of sampling to formally test the relative contributions of nosocomial versus community transmission, these findings suggest that colonisation prior to hospital admission may play a significant role in the pathogenesis of HAP. This observation is consistent with the suggestion that destabilisation of commensal communities could be a preceding indicator of infection^56^. Additional work, leveraging time-series samples, may shed further light on the role of early microbial perturbations and AMR carriage in the onset of HAP.

As many as 64% of samples harboured microbes that were predicted to have resistance to at least one antibiotic and ∼21% of isolates recovered were predicted to be multi-drug resistant based on putatively-acquired AMR gene content. These estimates (which will be on the low side, because they do not include resistance owing to mutation of chromosomal genes) far surpass the ∼13% prevalence estimated for culture-positive sepsis across 104 US hospitals^57^. Such a high rate of AMR gene carriage may reflect high antibiotic exposure in this patient demographic; with broad-spectrum antibiotics typically administered as soon as infection is suspected. Another contributing factor however could be the circulation of inherently-resistant bacterial strains that are already well-adapted to surviving in healthcare environments. Species such as *K. pneumoniae* and *P. aeruginosa*, are frequently implicated in nosocomial outbreaks and are among the most drug-resistant pathogens globally^58^. Our findings reiterate the importance of antimicrobial stewardship in managing HAP, and the potential utility of metagenomic sequencing to support this.

In conclusion, our work provides a comprehensive characterisation of the HAP microbiome, building knowledge apropos the wide diversity of organisms present. This may inform improved diagnostic strategies. Our analyses indicate that HAP and VAP are microbiologically overlapping conditions, with differences in AMR gene content of the microbial species identified but not in the distribution of species present. We speculate that factors such as host immune status and prior antimicrobial exposure may play a more central role in shaping microbial content than the specific context of infection. By reframing our understanding of VAP and recognising the broader array of organisms that may contribute to pneumonia, metagenomic approaches hold significant promise for enhancing diagnosis, guiding antimicrobial stewardship and improving patient outcomes.

## Methods

### Patient recruitment, samples and ethics

Surplus routine lower respiratory tract samples were included from patients at seven intensive care units (ICUs) participating in the INHALE WP1 study^26^ and processed at UCL, one of the two participating central laboratories for the study. Sputa, endotracheal tube aspirates and bronchoalveolar lavages were included if they had sufficient surplus volume (>1.4 ml) and were from patients in a participating ICU, hospitalised for ≥48 hours and about to receive a new antibiotic or change of antibiotic to treat new or worsening HAP or VAP. Samples for metagenomic analysis were eligible if they had been collected within 12 hours of antibiotic initiation, had been stored at 4°C, had not been frozen, and were depleted and extracted within 7 days of collection from the patient. Metagenomic analysis took place on samples that met these criteria from August 2017 onwards. Collection of clinical data (with consent or assent) included antimicrobial prescribing and clinical outcomes, and took place at four of the seven participating ICUs for a sub-set of participants.

Control group sputum was collected from eight healthy participants, recruited from Primary Care as a control group in a study approved by the London Hampstead Research Ethics Committee (14/LO/1409)^32^. These individuals self-identified as smokers, consistent with their ability to expectorate sputum for further analysis. At the time of sampling lung function parameters were normal, with participants free from acute respiratory illness. The INHALE WP1 study was approved by the UK Health Research Authority (Reference: 16/HRA/3882, IRAS ID: 201977) and the UCL DNA Infection Bank Committee, whose operation is governed by the London Fulham Research Ethics Committee (REC Reference: 17/LO/1530). Additional clinical details from a sub-set of WP1 patients were collected, with consent by ethical approval, from Camden and Kings Cross Research Ethics Committee, reference REC 16/LO/1618.

### Microbial culture

Standard-of-care microbial culture was carried out at the routine diagnostic laboratories serving each participating ICU according to standard operating procedures (SOPs). Laboratory SOPs followed the UK Standard for Microbiology Investigation of respiratory samples^59^. Briefly, each sample was homogenised with 0.1% dithiothreitol, followed by a 10^-5^ dilution, and inoculated onto a range of agars. In the case of BAL and NBL, culture was performed on serial dilutions of a sample that has first been concentrated by centrifugation. Plates are incubated for 40-48 hours, with identification of bacterial pathogens to species level by MALDI-TOF.

### PCR assays

Samples were tested at a centralised laboratory using both the Unyvero Pneumonia Panel (Curetis, Holzgerlingen, Germany) and the BioFire FilmArray Pneumonia Panel (BioFire Diagnostics, Salt Lake City, USA), both used according to manufacturer’s instructions^26^.

### Host depletion and DNA extraction

The sample processing and nanopore sequencing methods used have been published previously^22^. Briefly, respiratory samples that had been liquefied with Sputasol (Oxoid, UK) were treated with 2.2% saponin (Sigma, UK) for host cell lysis, followed by digestion with HL-SAN DNase (Articzymes, Norway). Samples were washed in PBS and centrifuged to pellet bacterial and fungal cells. The pellet was re-suspended in MagNA Pure Bacterial Lysis Buffer (Roche, Switzerland) and underwent bead-beating in a lysing Matrix E™ beads tube and a FastPrep 24 bead beater (MP Biomedical, USA) to disrupt microbial cell walls, followed by Proteinase K digestion. DNA was purified using the DiaSorin^®^ Arrow DNA Extraction Kit on the DiaSorin^®^ Ixt Extraction Platform, following manufacturer’s instructions, and eluted in 50 μL of elution buffer. DNA was quantified using the Qubit dsDNA HS assay kit for the Qubit 2.0 Fluorometer (Thermo Fisher Scientific, USA). Extracts were further cleaned and concentrated using AMPure XP Beads (Agencourt, France). All batches of samples were processed along with a PBS negative control through the whole process.

### MinION library preparation and sequencing

Library preparation was performed using the Rapid PCR Barcoding Kit RPB004 (ONT) with a modified protocol to amplify very low DNA inputs as previously described^22^. Following PCR, DNA was quantified using the Qubit dsDNA HS kit and pooled in equal concentrations with 6 samples per run plus a negative process control. Libraries were SPRI cleaned with AMPure XP beads and analysed using the Qubit dsDNA BR kit and the Agilent Bioanalyzer 2100 using the DNA 12000 kit (Agilent Technologies, USA). The library was loaded onto nanopore flow cells (R9.4.1) with sequencing performed on the MinION platform (ONT).

### Sequencing data pre-processing and quality control

Basecalling of nanopore sequencing data, followed by adapter, primer and barcode trimming, was performed using Guppy v1.1. Human reads were removed by aligning all reads to the human T2T-CHM13v2.0 reference genome (GCF_009914755.1) with minimap2 v2.26^31^ (‘map-ont’ preset).

### Taxonomic classification of sequencing reads

Taxonomic classification of human-filtered sequencing reads was performed using Kraken2 v2.1.2^60^ against the Plus PFP database curated by Ben Langmead (https://benlangmead.github.io/aws-indexes/k2; updated 12 Jan 2024), which comprises archaea, bacteria, viral, plasmid, human, protozoa and plants reference sequences. Unclassified reads and reads classified as human or plant were removed from further taxonomic analysis. We then calculated the relative abundance of a taxon in each sample as the number of reads divided by the total number of remaining reads in the sample. To minimise the effects of taxonomic misclassification and sequencing artefacts, taxonomic assignments where the relative abundance was 0.5% or less, or where no more than 10 reads were assigned were considered false positives and were removed from further analysis. The reads assigned to these taxa were set to zero, and the relative abundance of remaining taxa were re-calculated.

To minimise the effects of laboratory contamination, we considered removing all taxa that were found in the negative sequencing controls. However, the most prevalent HAP pathogens (e.g. *E. coli, S. aureus, P. aeruginosa*) were also commonly found in these negative controls. This strict decontamination approach would therefore remove many potentially genuine signals. We therefore opted for a less stringent decontamination heuristic, where we removed taxa whose relative abundance was less than twice the relative abundance of those in negative sequencing controls of the same run (also see **Supplementary Note 1**).

### De novo metagenomic assembly

Human-filtered sequencing reads were assembled using metaFlye v2.9.4^61^ and contigs were polished using Medaka v1.11.3 (https://github.com/nanoporetech/medaka.git). Taxonomic binning of contigs to produce draft assemblies was performed using metaBat v2.15^62^. Taxonomic classification and quality assessment of taxonomic bins were performed using GTDB-tk v2.4.0^63^ and CheckM2 v1.0.1, respectively. Taxonomic bins that could not be classified at the species level, or which contained gene markers spanning multiple domains, were removed.

### AMR phenotypic prediction

AMR resistance phenotypes for all samples were predicted based on AMR gene content using ResFinder v4.1.11^41^. Predictions were made using either sequencing reads (in the *fastq* format), or using de novo assembled taxonomic bins (in the *fasta* format) as input. For the former, we specified the ‘--nanopore’ flag – which enables the use of optimised alignment parameters for error-prone nanopore sequence data – a minimum gene coverage of 80% and a minimum sequence identity of 90%. Additionally, to minimise the effects of laboratory contamination, we excluded phenotypic predictions if the same predictions were made for negative sequencing controls in the same run. For the latter, as some genes may be split across more than one contig due to fragmented assembly, we relaxed the minimum gene coverage threshold to 50%, though this had minimal impacts on our results. Chromosomal genes conferring intrinsic resistance were removed, with the list of exclusions provided in **Supplementary Table 5**. The classification of antibiotics into antibiotic classes are provided in **Supplementary Table 6**.

### Phylogenetic analysis

For each of the top three species for which we recovered the highest number of complete genome assemblies (i.e., *S. aureus*, *H. influenzae* and *E. coli*), we compiled separate datasets of publicly-available bacterial assemblies from the AllTheBacteria v0.2^64^ database. For *S. aureus* and *E. coli*, we retrieved all human-associated sequenced isolates collected in the United Kingdom. Due to the relatively less intensive sampling of *H. influenzae* in the UK, we considered isolates collected worldwide. For each set of sequences, we reconstructed initial neighbour-joining trees using Mashtree v1.4.6^65^, which leverages the alignment-free, k-mer-based, Mash distances^66^ between sequences. Using these initial trees, we then performed phylogenetically-informed subsampling of the datasets using Treemmer v0.3^67^ to reduce genetic redundancy. We specified the ‘-RTL 0.90’ flag, subsampling each dataset to 90% of the initial genetic diversity represented. For these subsampled datasets, we then reconstructed more robust, maximum-likelihood phylogenies using a core-genome alignment approach. To do so, we first annotated each assembly using Prokka v1.14.6^68^ and generated a concatenated core-gene codon alignment using Panaroo v1.5^69^. We extracted the variable sites in these alignments using snp-sites v2.5.1^70^, and removed sequences and sites that had >20% missingness. We then performed maximum-likelihood phylogenetic reconstruction on these SNP alignments under a GTR+Γ model using VeryFastTree v4.0.3^71^. To minimise the effects of recombination, we pruned putatively recombinant sites using Gubbins v3.3.1^72^ before re-estimating recombination-pruned maximum-likelihood phylogenies. Note that since we are using a SNP-alignment, by definition, all invariant sites have been removed, but these are important for accurate estimation of branch lengths. While the topologies shown in **Fig. 5a-c** are not affected by this, we caution against interpreting the branch lengths shown.

To compute the pairwise SNP distances shown in **Fig. 5e**, we used the dist.dna function in the R package, Ape v5.7.1^73^, on the concatenated core-genome codon alignments. The expected pairwise SNPs were estimated using the following formula:

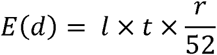

Where *d*, *l*, *t*, and *r*, denote the number of pairwise SNPs, alignment length, time difference in weeks, and assumed substitution rate in substitutions/site/year, respectively.

### Data analysis and visualisation

All data analyses were performed using R v4.3.1. All visualisations were performed using *ggplot* v3.4.2^74^ or *ggtree* v3.8.2^75^.

## Acknowledgments

This research was funded by the National Institute for Health Research (NIHR) under its Programme Grants for Applied Research Programme (Reference Number: RP-PG-0514-20018). C.C.S.T. is funded by the National Science Scholarship from the Agency for Science, Technology and Research (A*STAR), Singapore. L.v.D. is funded by a UKRI Future Leaders Fellowship (MR/X034828/1). F.B. and additionally L.v.D are funded by the European Commission (Horizon 2021-2024, END-VOC Project). Infrastructure support for DB was provided by the NIHR University College London and Biomedical Research Centres. The views expressed are those of the authors and not necessarily those of the NIHR, or the Department of Health and Social Care. Views and opinions expressed are however those of the authors only and do not necessarily reflect those of the European Union or the European Health and Digital Executive Agency. For the purpose of open access, the corresponding author has applied a ‘Creative Commons Attribution’ (CC BY) licence to any Author Accepted Manuscript version arising. The authors acknowledge the use of the UCL Computer Science cluster and associated support services, in the completion of this work.

## Author contributions

S.R, A. A. and D. O. collected samples and laboratory data and performed metagenomic sequencing. A. A. and D. O. generated PCR data. A. A. and T. C optimised the metagenomic sequencing workflow. DB, MP and JH recruited patients, provided clinical input and collected data. C.C.S.T performed all bioinformatic and statistical analyses. C.C.S.T., L.v.D., F.B., and V.E. wrote the manuscript with intellectual inputs from all co-authors. VE, VG, JOG and DML obtained funding. V.E., J.O.G., DB, C.C.S.T, L.v.D. and F.B. conceived the work. J.O.G., V.E. and T.McH. supervised the laboratory work.

## Ethics approval

This study was approved by UK Health Research Authority (Reference: 16/HRA/3882, IRAS ID: 201977) and the UCL DNA Infection Bank Committee, whose operation is governed by the London Fulham Research Ethics Committee (REC Reference: 17/LO/1530).

## Data and code availability

All custom code used to perform the analyses reported here are available on GitHub (https://github.com/cednotsed/pneumonia).

## Extended Data Figures

**Extended Data Figure 1.**
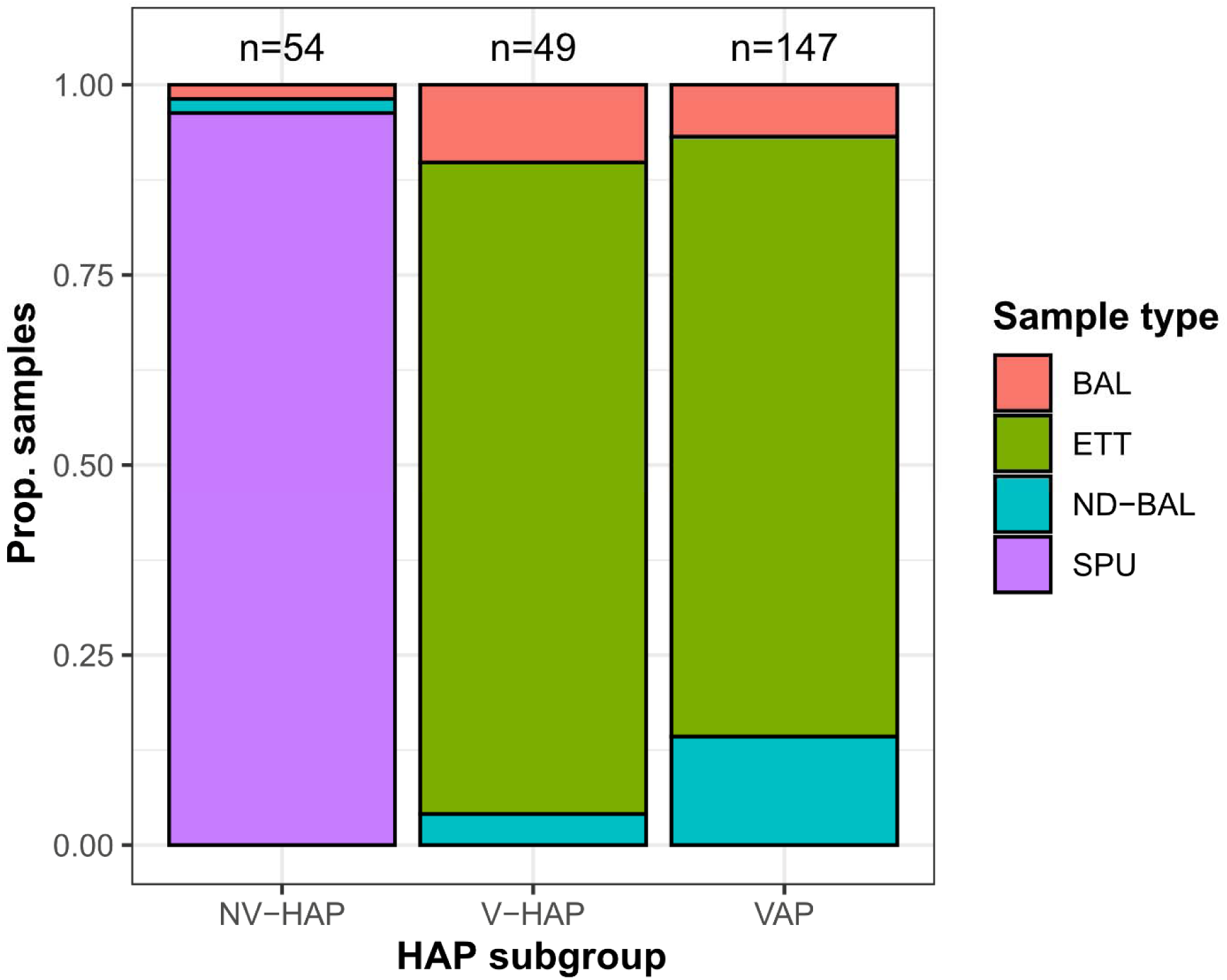
Summary of sample types represented in this dataset stratified by HAP subgroup.

**Extended Data Figure 2.**
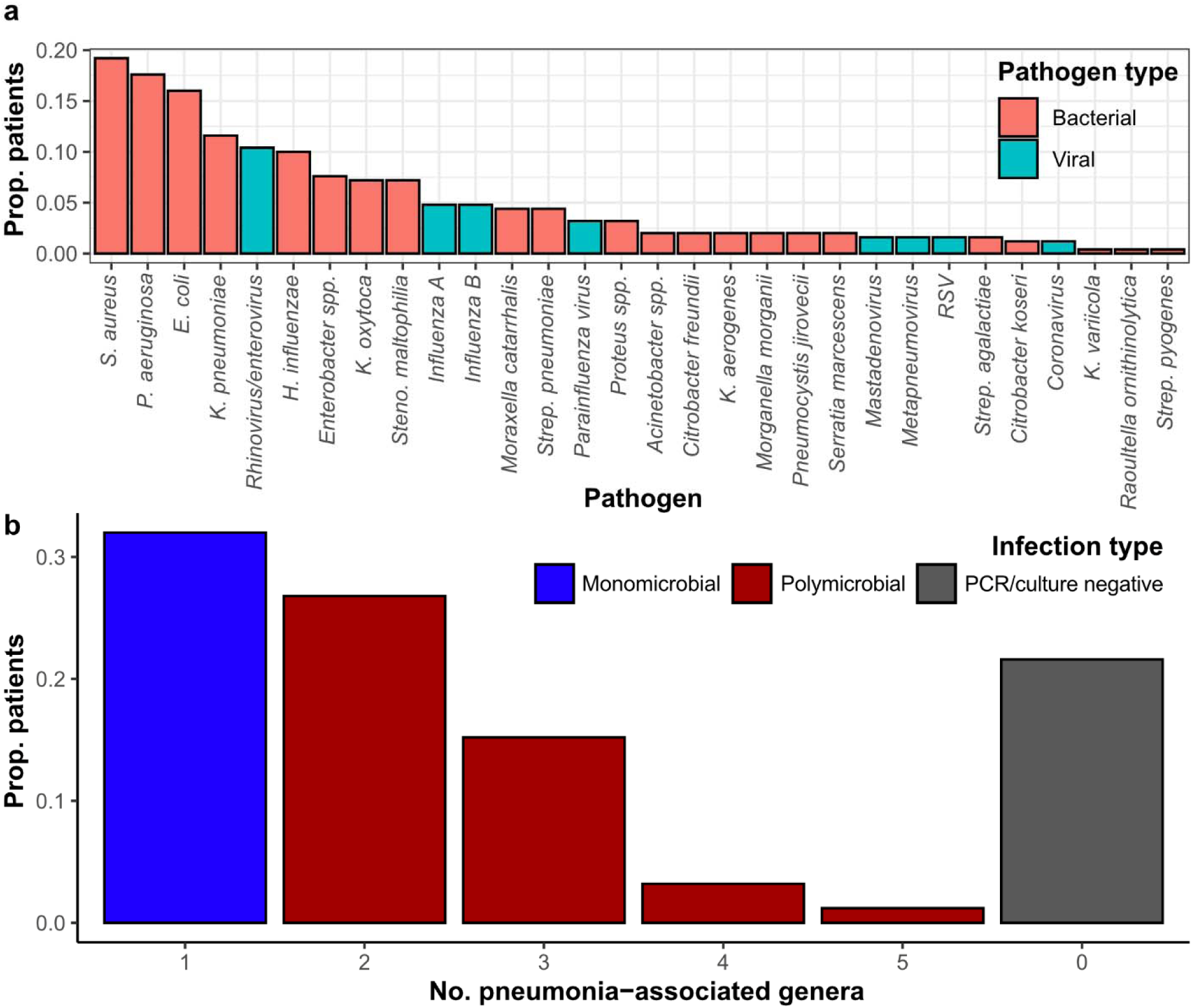
Microbial characteristics of samples as assessed by PCR/culture. (a) Proportion of patients (n=250) testing positive for each of the 30 most common pathogens detected through PCR and/or culture. (b) Barplot providing the number of infections found to harbour pneumonia-associated genera. Polymicrobial infections are defined by the detection of more than one pneumonia-associated genus by PCR and/or culture.

**Extended Data Figure 3.**
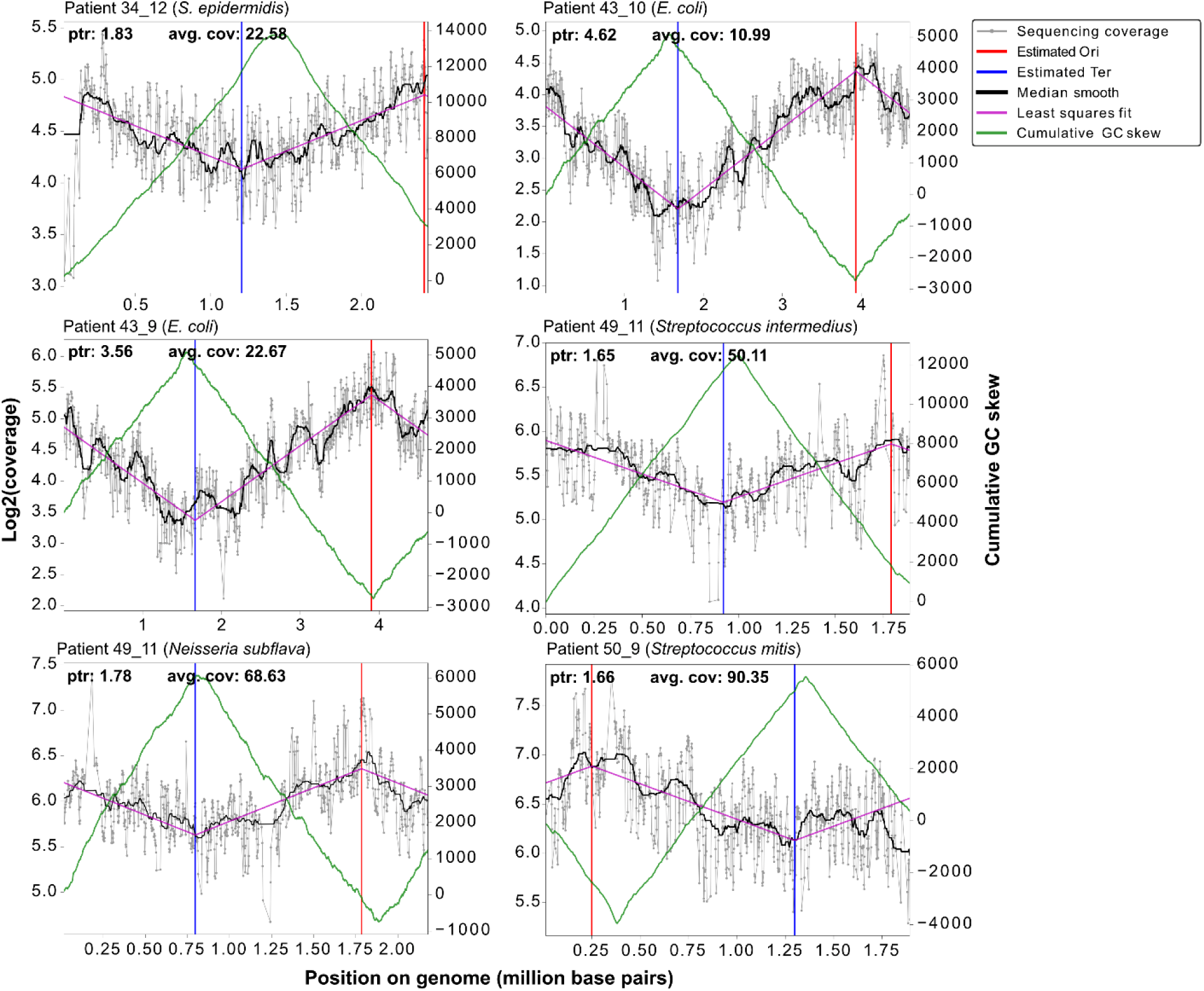
Molecular signatures of replicating bacteria in PCR/culture-negative HAP patients. Replication rate analyses^29^ were performed for all bacterial species that were assigned at least 10,000 reads by Kraken2 in the 49 HAP samples that were PCR/culture negative. Six bacterial species in five patient samples showed sequencing coverage patterns characteristic of replicating bacterial populations in microbial cultures^28^ (see Methods). In growing bacterial populations, sequencing coverage is higher nearer to the origin of replication (Ori) and lower nearer to the terminus (Ter) of the bacterial chromosome. Each plot shows this characteristic sinusoidal shape of sequencing coverage across the bacterial genome for each of these species. The Ori and Ter positions inferred from these coverage plots were concordant with those inferred using an orthogonal method based on GC skew across the genome^29,76^.

**Extended Data Figure 4.**
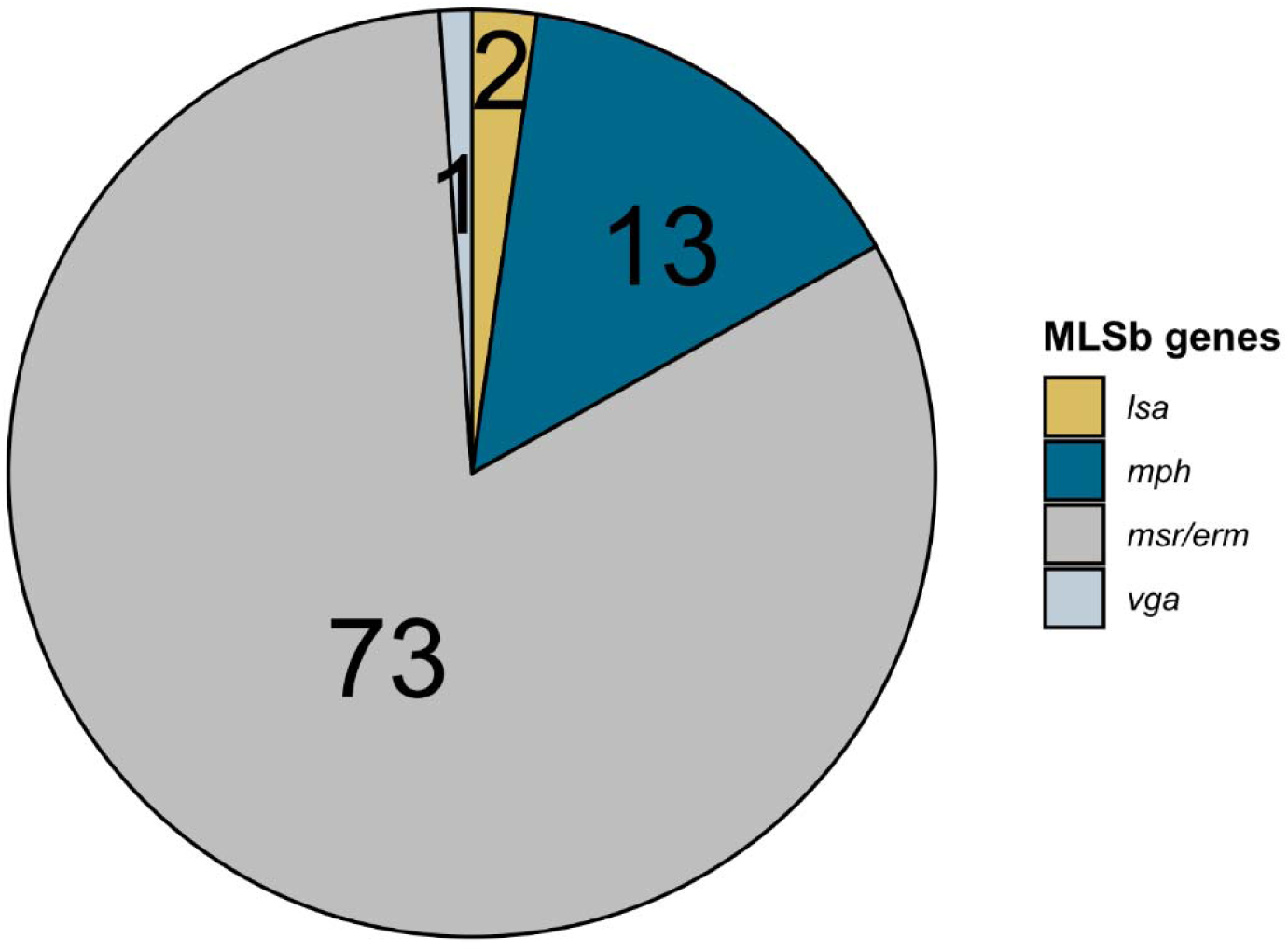
Frequency of detected AMR genes associated with resistance to some or all macrolide, lincosamide or streptogramin B antibiotics, as determined by ResFinder^41^ across the dataset.

**Extended Data Figure 5.**
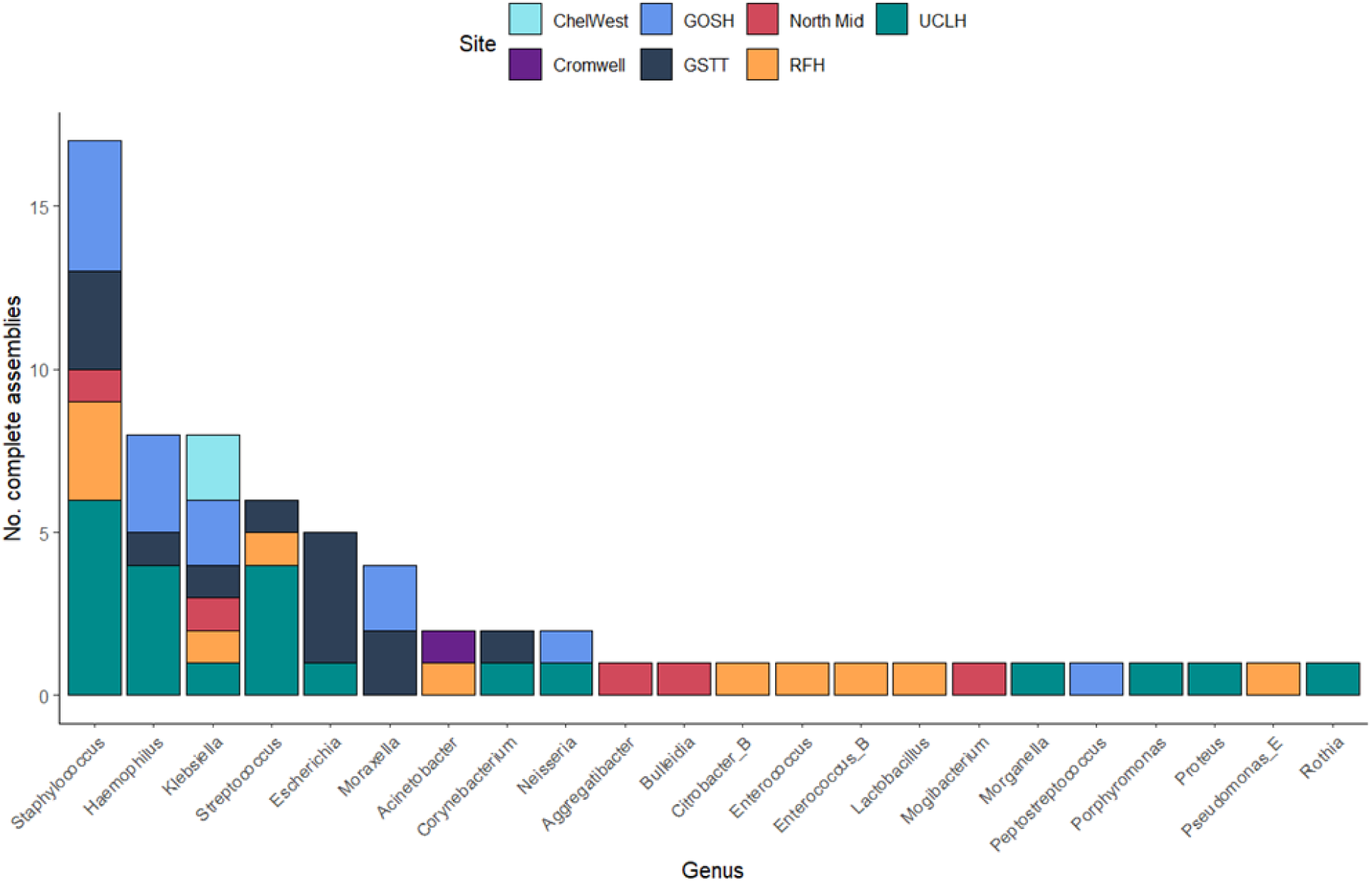
Summary of high-quality complete bacterial assemblies recovered. Barplot providing the number of bacterial assemblies assessed by CheckM2^45^ to be at least 90% complete and of high quality, stratified by bacterial genus. The 66 assemblies recovered were assigned GTDB-harmonised genus names using GTDB-tk^77^.

**Extended Data Figure 6.**
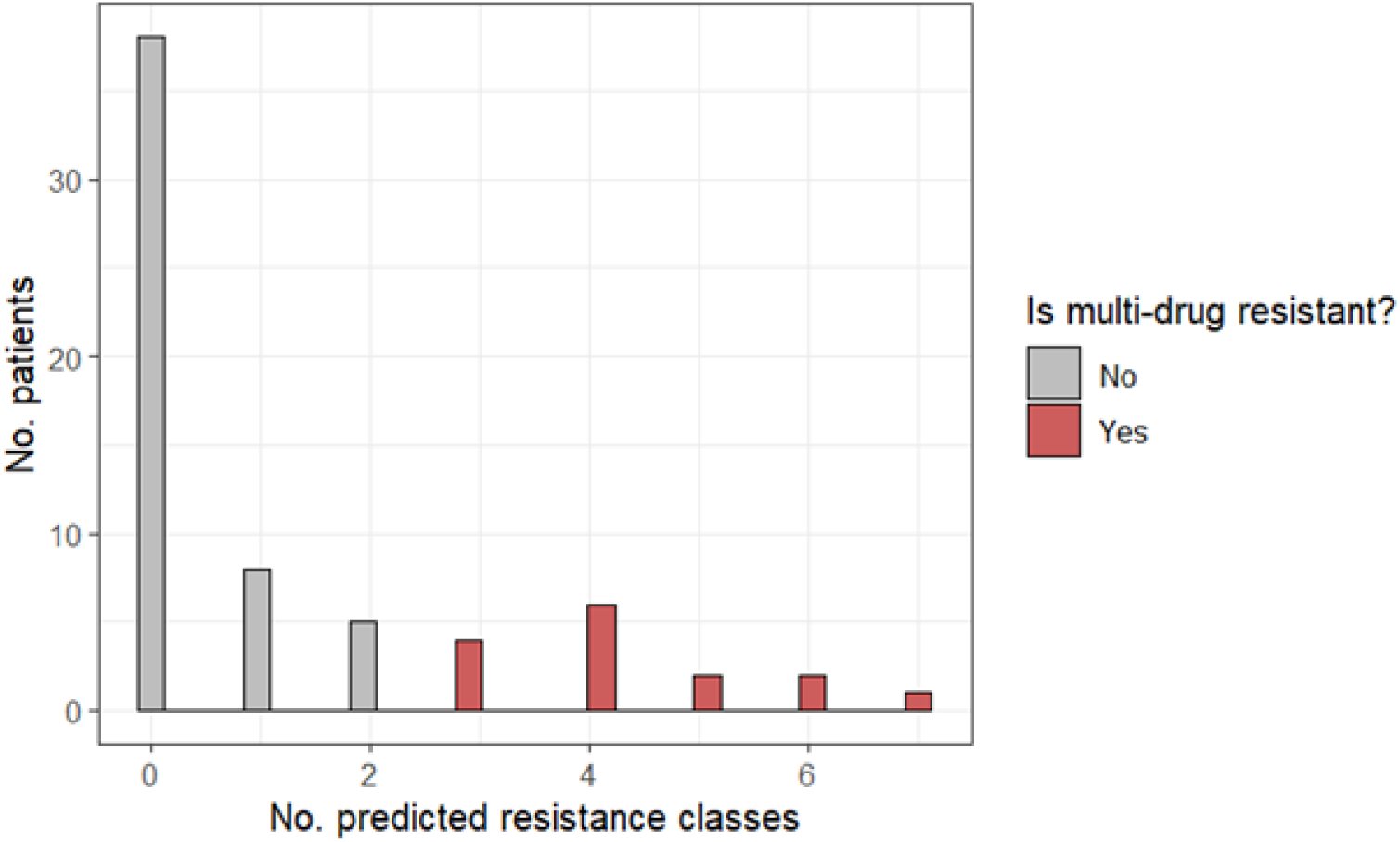
High prevalence of predicted drug resistance in recovered bacterial assemblies. Histogram showing the number of predicted AMR resistance classes recovered per patient sample. Chromosomal genes conferring intrinsic resistance were removed (**Supplementary Table 5**). The antibiotic classes considered are listed in **Supplementary Table 6**.

