## Supplementary Note 1 for "Characterising the microbial and antimicrobial resistance signatures of hospital-acquired pneumonia using nanopore metagenomic sequencing"

Prior to our metagenomic analyses, we applied a set of bioinformatic filters to reduce noise due to index hopping, taxonomic misclassification and laboratory contamination. In the Kraken2 taxonomic profiles of all samples, taxonomic assignments whose associated relative abundance was at most 0.5% or with at most 10 reads assigned were considered false positives and removed from further analysis. This, in principle, minimises the effects of index hopping, where sequencing reads from one sample are erroneously assigned barcodes from another one within the same run during demultiplexing. The rate of barcode crosstalk for the sequencing flowcell version we used (R9.4.1) was estimated previously to be around 0.056%[1] so our relative abundance threshold of 0.5% is likely stringent enough. To test this empirically, we compared the total number of reads per taxon across all patient samples and of the negative control in each run. A high correlation would indicate that barcode crosstalk is a significant confounder in our characterisation of microbial profiles. However, the correlation between the taxon abundance in samples and controls was extremely weak (Pearson's $r$=0.068), suggesting that the effects of barcode crosstalk are minimal.

The use of negative controls for each sequencing batch enabled us to account for laboratory contamination. Our decontamination approach involved removing taxa whose relative abundance was less than 2x the relative abundance of those in negative sequencing controls of the same run. The number of microbial reads associated with the negative controls was generally much lower than that for patient samples (median reads=1665 and 75,608, respectively). However, there were eight sequencing batches where the negative controls had >10,000 microbial reads. Inspection of the microbial profiles in these sequencing batches prior to and following decontamination indicated that our approach effectively removed contaminants while retaining biologically relevant taxa (**Supplementary Fig. 1**). Additionally, the mean Bray-Curtis distance between patient samples and the negative controls within runs increased from a mean of 0.759 to 0.959 after decontamination, indicating that the microbial profiles of patient samples became more distinct from those of the negative controls after filtering (**Supplementary Fig. 2**). Separately, we tested whether the choice of decontamination threshold affects our results by repeating some of the analyses reported in the main text using various decontamination thresholds (1x, 2x, 5x and 10x). Across all thresholds, the microbial profiles of the different HAP subtypes could not be clearly separated (**Supplementary Fig. 3**), suggesting that these results are robust to the choice of decontamination threshold.

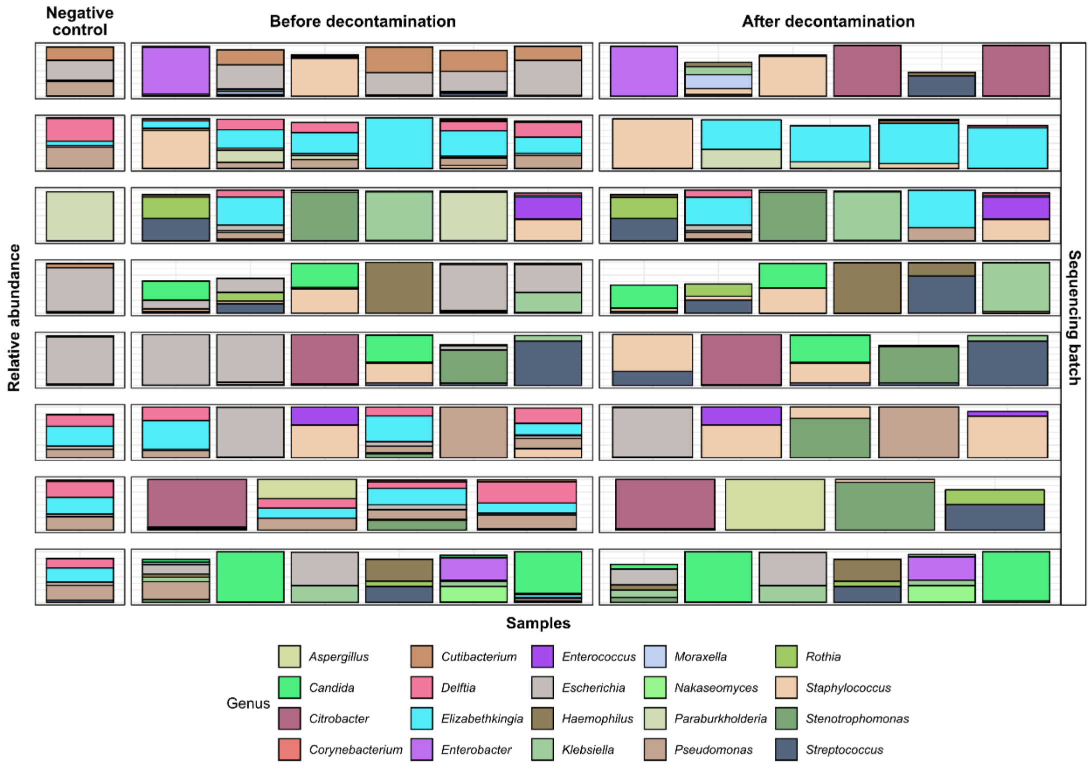


***Supplementary Figure 1.*** *Relative abundance of negative controls and patient samples in sequencing batches whose negative controls had a high microbial read count (>10000 reads). The microbial profiles of patient samples before and after decontamination was applied are shown. For simplicity, we only show the relative abundance of the top 20 most abundant taxa, as assessed by mean relative abundance across all patient samples.*

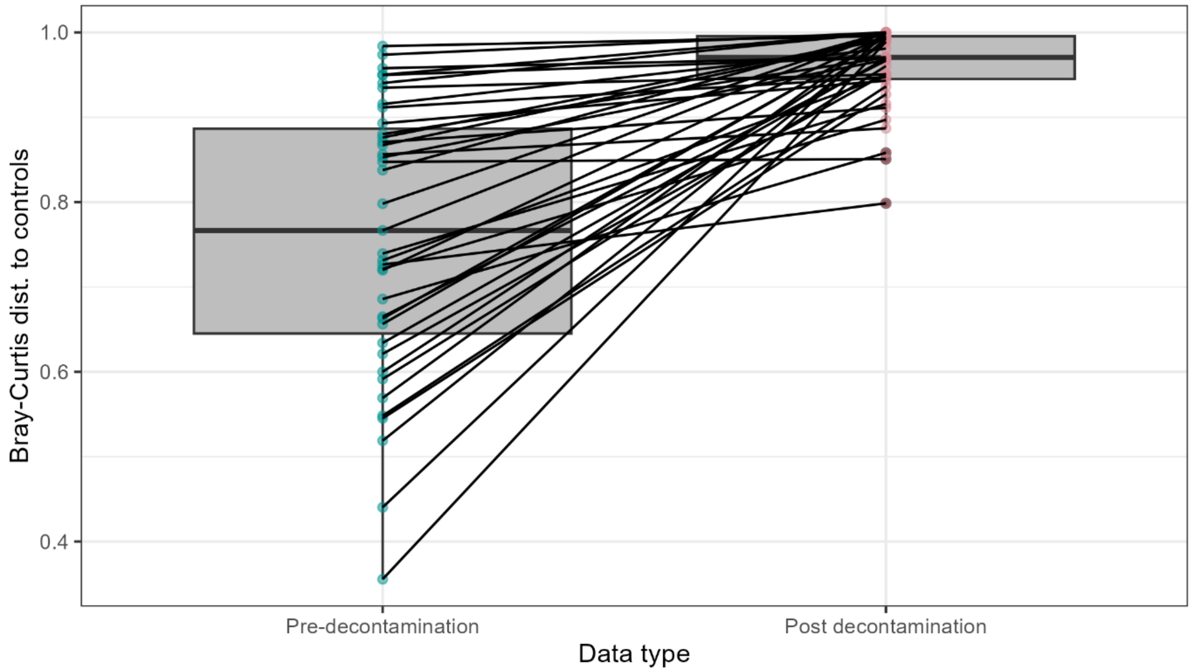


***Supplementary Figure 2.*** *Bray-Curtis distance of patient samples from the corresponding negative controls in the same run before and after decontamination.* Boxplot elements are defined as follows: centre line, median; box limits, upper and lower quartiles; whiskers, 1.5x interquartile range.

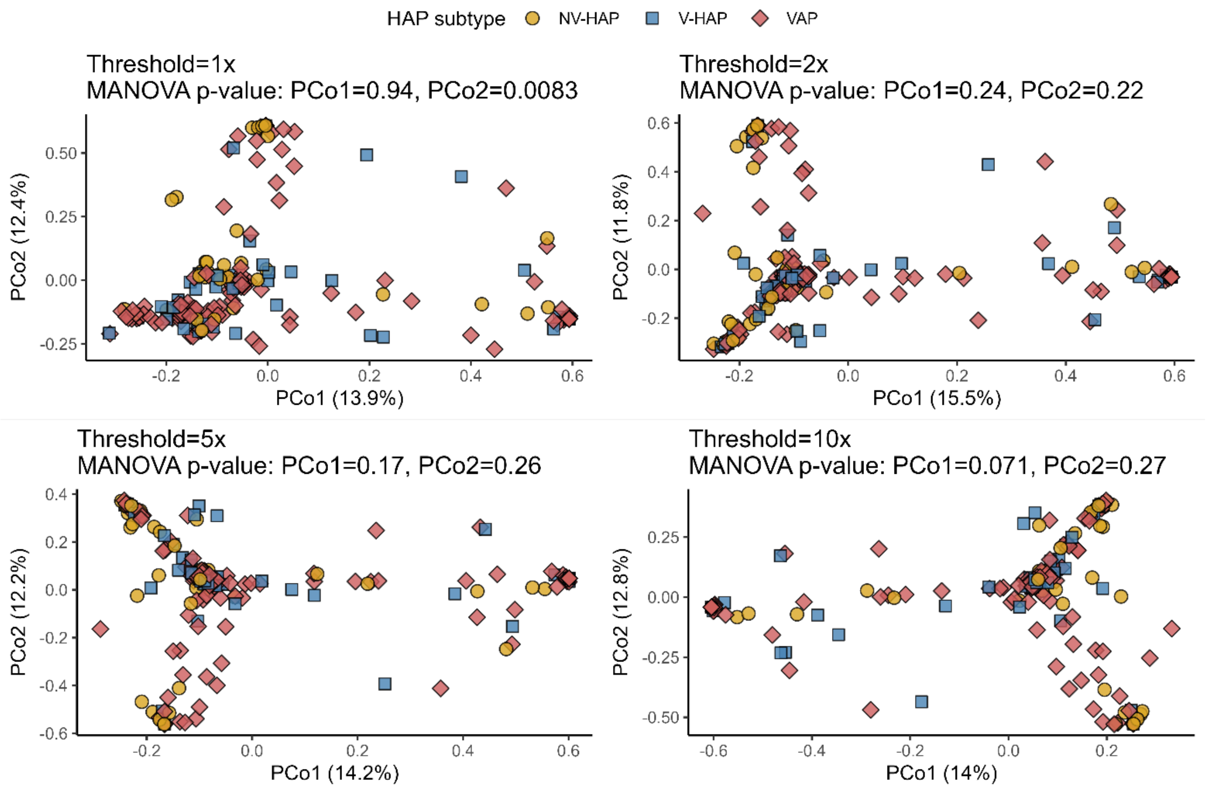


**Supplementary Figure 3. Analysis is robust to choice of decontamination threshold.** Principal coordinates analysis of Bray-Curtis distances, and boxplots showing the lack of clustering in the microbial profiles of NV-HAP, V-HAP and VAP samples (as in **Fig 3a**). The PCoA analysis was performed using various decontamination thresholds (1x, 2x, 5x, and 10x). The p-values of the MANOVA test for each principal coordinate (PCo) – which corrects for the effects of sequencing depth and sampling site – are annotated.
